# Acute Sublethal Heat Stress Impairs Blood Feeding and Trypanosome Infection in the Kissing Bug, *Rhodnius prolixus*

**DOI:** 10.64898/2026.01.27.701963

**Authors:** Syeda Farjana Hoque, Paige Crawford, Ann Miller, Joshua Tompkin, Milad Ahmed, Asima Das, César González Zermeño, Noelia Lander, Joshua B. Benoit

## Abstract

Kissing bugs are the primary vectors of *Trypanosoma cruzi*, the causative agent of Chagas disease. Kissing bugs are exposed to thermal variability, including short periods of heat stress, which can induce mortality or exert sublethal effects. This study investigated *Rhodnius prolixus* following brief periods of high thermal stress with respect to survival, blood feeding, developmental processes, and *T. cruzi* infection, with a focus on sublethal effects. Our results demonstrated a significant decrease in survival for *R. prolixus* at 42 °C for 8 hours. When exposed to sub-lethal thermal stress (40°C for 8 hours), blood ingestion (amount and proportion) was reduced after 24 hours of recovery from thermal stress. Among the bugs that fed after 24 hours, molting was not impacted by temperature exposure. The infection rate decreased after heat exposure, likely due to reduced blood volume ingested when feeding 24 hours after heat stress. A week of recovery after exposure to higher temperatures improved feeding and increased infection rates to levels comparable to those of kissing bugs not exposed to thermal stress. Our findings offer insights into how extreme temperature events may influence Chagas disease. Specifically, these studies highlight the need to clarify how temperature, particularly at sublethal levels, interacts with vector biology to alter parasite transmission.

## Introduction

Kissing bugs (Hemiptera: Reduviidae) are obligate blood feeding insects and the primary vectors of *Trypanosoma cruzi*, the parasite that causes Chagas disease. Chagas disease is endemic in twenty-one countries in Latin America (Nunes et al., 2018). Recent evidence suggests that Chagas disease is endemic in the southern United States, with infections identified across several southern states, likely due to local transmission (Beatty et al., 2025; Farani et al., 2025). According to the World Health Organization (WHO), around 6-8 million people worldwide are infected with *T. cruzi*, and around 70 million are at risk of infection (Lidani et al., 2019). Kissing bugs can live in various thermal environments, ranging from sylvatic habitats, such as palm tree crowns and bird nests, which are relatively thermally buffered, to domestic settings, including thatched and adobe houses, where daily temperature fluctuations and acute heat extremes are more pronounced. (Lorenzo and Lazzari, 1999; Sanchez-Martin et al., 2006; Abad-Franch et al., 2015). This exposes kissing bugs to highly variable environmental conditions, particularly temperature, which likely plays a central role in shaping their biology and vectorial capacity.

Thermal changes affect the survivability, blood feeding, host-seeking behavior, molting, and disease transmission in kissing bugs (Settembrini, 1984; Guarneri et al., 2003). Daily temperature cycles have been shown to entrain circadian rhythms and influence behavioral activity of triatomines (Fresquet and Lazzari, 2014; Steel and Vafopoulou), as well as through other non-photic cues (Valentinuzzi et al., 2014). Like many ectothermic animals, triatomines exhibit behavioral thermoregulation, adjusting their location based on thermal preferences and ecological context (Vielmetter, 1958; Remmert, 1960; Wurtsbaugh and Neverman, 1988; van Dijk et al., 2002). Conversely, exposure to high temperatures is associated with greater mortality, enhanced rates of water loss, and depletion of nutritional resources (Beament, 1961; Okasha, 1968; Loshouarn and Guarneri, 2024). Overall, the thermal shifts have direct implications for the biology of kissing bugs. Temperature plays a critical role in shaping interactions between kissing bugs and *T. cruzi*. Higher temperatures can accelerate parasite development, increase multiplication rates, and reduce the time to parasite appearance in vector feces, potentially enhancing transmission efficiency (Wood, 1954; Phillips, 1960; Kollien and Schaub, 2000; Elliot et al., 2015; Loshouarn and Guarneri, 2024). At the same time, thermal stress can amplify the detrimental effects of infection on the vector, including increased mortality and delayed molting (Botto-Mahan, 2009; Elliot et al., 2015; Loshouarn and Guarneri, 2024).

Previous studies have focused on how sustained increases in temperature or long-term exposure to fluctuating thermal environments alter triatomine development, feeding performance, and *T. cruzi* development (Rolandi and Schilman, 2018; Tamayo et al., 2018; Álvarez-Duhart et al., 2024). In contrast, the effects of brief, sublethal acute heat stress followed by recovery on feeding and parasite acquisition remain poorly characterized. Short-term thermal stress events are likely to become more frequent under climate change (Perkins-Kirkpatrick and Lewis, 2020; Marx et al., 2021), underscoring the need to understand their consequences for vector–parasite interactions. In this study, we evaluated the impact of acute heat stress on kissing bug biology. To do so, we assessed the survival of *R. prolixus* across a temperature gradient (24-44 °C) to establish lethal and sublethal thermal exposures. Following the assessment of survival, we examined how sublethal heat stress (32 °C & 40 °C) affects blood consumption, survival, and molting rates at one and seven days after heat stress. Lastly, we assessed the level of trypanosome infection following sublethal heat stress (40 °C) in relation to blood ingestion. These studies provide evidence that short periods of sublethal heat stress directly affect aspects of kissing bugs that are likely to reduce trypanosome transmission.

## Materials and Methods

### Kissing bug rearing

Kissing bugs (*R. prolixus*) were obtained from the colonies established at the Centers for Disease Control and Prevention (BEI Resources, NR-44077) (Wormington et al., 2018). The colony was maintained at 24 ± 1 °C, 50 ± 10% relative humidity (RH), and a 12:12 h light: dark cycle. Bugs were fed defibrinated rabbit blood using an artificial feeder (Hemotek® PS5 feeder, 37 °C, Parafilm® membrane) every two weeks (Ahmed et al., 2025; Carlson et al., 2025 May 23). Fourth-instar nymphs (4-5 weeks unfed) were used in heat exposure and infection assays.

### Heat stress exposure

Heat treatments were conducted using a Fisher Scientific Isotemp Digital Dry Block Heater (Model 2001, Cat. No. 11-715-125D) according to previously developed methods (Chakraborty, Zigmond, et al., 2025; Chakraborty, Shah, et al., 2025). Each bug was placed in a plastic vial (≈26 mL capacity) containing small strips of filter paper and brown paper to provide the bugs with a proper environment. The vial lids were modified to feature a mesh opening for airflow. Plastic vials were placed in the metal block wells of the dry bath. The entire dry block was covered to ensure a consistent temperature in the vials. To assess sub-lethal heat tolerance, nymphs were exposed to 24, 28, 32, 36, 40, or 44 °C for 8 h. For each temperature, 36 insects were tested (six groups of six), yielding a total of 216 individuals across all treatments. Survival was assessed by counting live and dead individuals based on responsiveness to gentle stimulation immediately after heat exposure and at 1 and 7 days post-exposure (Figure 1). Temperature was verified at the start of each run with a calibrated thermocouple (±0.2 °C, Omega). After exposure, all insects were returned to standard rearing conditions (24 °C, 60% RH).

**Figure 1.**
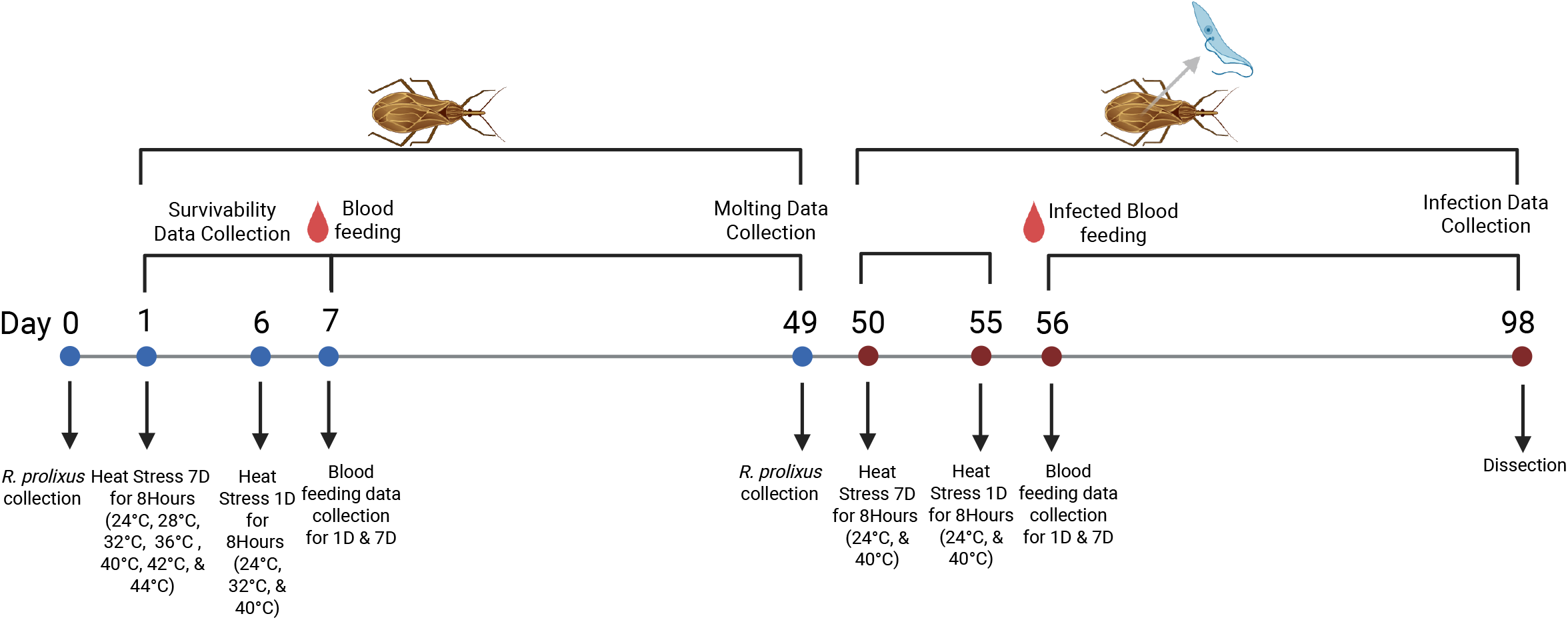
Summary of the experimental design used to evaluate the effects of acute heat stress on Rhodnius prolixus. Fourth-stage unfed nymphs were collected and subjected to heat-stress treatments for 8 hours at 24–44 °C (n = 36 per temperature). The survivability of kissing bugs was assessed under normal colony conditions and at lethal temperatures (24 °C, 28 °C, 32 °C, 36 °C, 40 °C, 42 °C, and 44 °C). Blood feeding and molting results were collected in 1-day (1D) and 7-day (7D) recovery groups for control (24 °C), medium point (32 °C), and sub-lethal point (40 °C). Separate cohorts were infected with *Trypanosoma cruzi*, and the effects of infection rate and recovery period on their infection rate and recovery period were measured at 24 °C and 40 °C.

### Feeding and molting assays

To determine the amount of blood ingested, feeding success, and molting success, bugs were exposed to 24 °C, 32 °C, and 40 °C (sublethal heat) for 8 hours. At 1 day and 7 days after heat exposure, bugs were weighed (±0.1 mg) and offered defibrinated rabbit blood (37°C) for 30 minutes. Post-feeding mass was recorded immediately after feeding to determine the amount of blood ingested. Nymphs were then maintained at 24 °C, 50 ± 10% RH, and monitored daily for ecdysis. For the feeding experiment, we used 216 insects at three temperature treatments.

### Infection assays with *T. cruzi* parasite

Wild-type *T. cruzi* epimastigotes were cultured at 28 °C in liver infusion tryptose (LIT) medium supplemented with 10% heat-inactivated fetal bovine serum (FBS), penicillin (100 IU/mL), and streptomycin (100 µg/mL) (Bone and Steinert, 1956). Cultured epimastigotes in the exponential growth phase (1-2 × 10^7^ cells/mL) were used for triatomine infection. Fourth-instar *R. prolixus* were exposed to either 4 °C or 24 °C (control) for 8 h. At 1 and 7 days post-temperature exposure, insects were weighed and fed blood meals containing *T. cruzi* epimastigotes. Briefly, parasites were washed in 5 mL of sterile 1× PBS (pH 7.4) and mixed with defibrinated rabbit blood that had been complement-inactivated at 56 ± 0.5 °C for 45 minutes prior to parasite addition. Blood meals were provided using an artificial feeder maintained at 40 °C, with a final density of 1 × 10^8^ parasites/mL (Fellet et al., 2014; Vieira et al., 2018; Batista et al., 2020). A total of 144 insects were used in this study across the two temperature conditions and two post-exposure time points (1 and 7 days). The kissing bugs were maintained at 24 ± 0.5°C, 50 ± 10% relative humidity, and a 6:00 am/6:00 pm light/dark photoperiod. Four weeks after infection, hindguts were dissected, emulsified and homogenized in 100 µL of sterile 1× PBS (pH 7.4). Parasite presence was assessed under a light microscope to determine the percentage of infected insects (Ahmed et al., 2025; Carlson et al., 2025 May 23). Infection status was evaluated using six to nine independent groups, each consisting of 3-6 infected insects.

### Statistical analysis

Survival curve data were analyzed using R (version 4.3.2). For each temperature point, the percentage of surviving insects was calculated by dividing the number of surviving individuals by the total number exposed and multiplying by 100. A nonlinear regression model was fitted to the data using nls () function in base R (The R Project for Statistical Computing). The model followed a sigmoidal dose–response function (Motulsky and Christopoulos, 2004).

Survival (%) = 100 / (1 + e^(−a(T − b))), where **T** is the Temperature, **a** is the slope of the curve, and **b** corresponds to the estimated lethal dose at which 50% of the individuals died (LD_50_). Model fitting was performed using least-squares estimation, and initial parameter values were chosen empirically. The LD_50_ was extracted directly from the fitted model coefficients. Predicted survival values were calculated across a continuous temperature range (24–44 °C), and the model fit was visualized alongside the observed survival percentages. Horizontal and vertical dashed lines were used to denote the 50% survival threshold and the corresponding LD_50_ estimate, respectively.

Blood meal size, feeding proportion, and molting were analyzed across temperature treatments using two-way ANOVA to assess the effects of temperature and recovery period. When significant main effects or interactions were identified, Tukey’s multiple comparisons test was used to evaluate pairwise differences among groups. A threshold of p <= 0.05 was considered statistically significant. All data are presented as mean +/-SEM, and statistically significant differences are shown in the figures using significance markers (for example: *, **, ***).

Statistical analyses for these experiments were conducted using GraphPad Prism 10.4.1 (GraphPad Software, San Diego, CA, USA), except for infection status, which was assessed using a generalized linear model (binomial logit) in R.

## Results

### Survival of *R. prolixus during acute heat stress*

Survival of *Rhodnius prolixus* fourth-instar nymphs declined sharply with increasing temperature following an 8-hour heat exposure (Figure 2). All the kissing bugs survived at 24 °C, with the survival rate remaining at 40 °C (78%). The survival rate declined at 42 °C, with only 28% of kissing bugs surviving. Mortality was 100% at 44 °C. A logistic regression model estimated the lethal dose for 50% mortality at 41.1 °C (Figure 2). These results indicate a narrow thermal threshold beyond 40 °C, above which survival rapidly declines, highlighting the sensitivity of *R. prolixus* to acute high-temperature stress.

**Figure 2.**
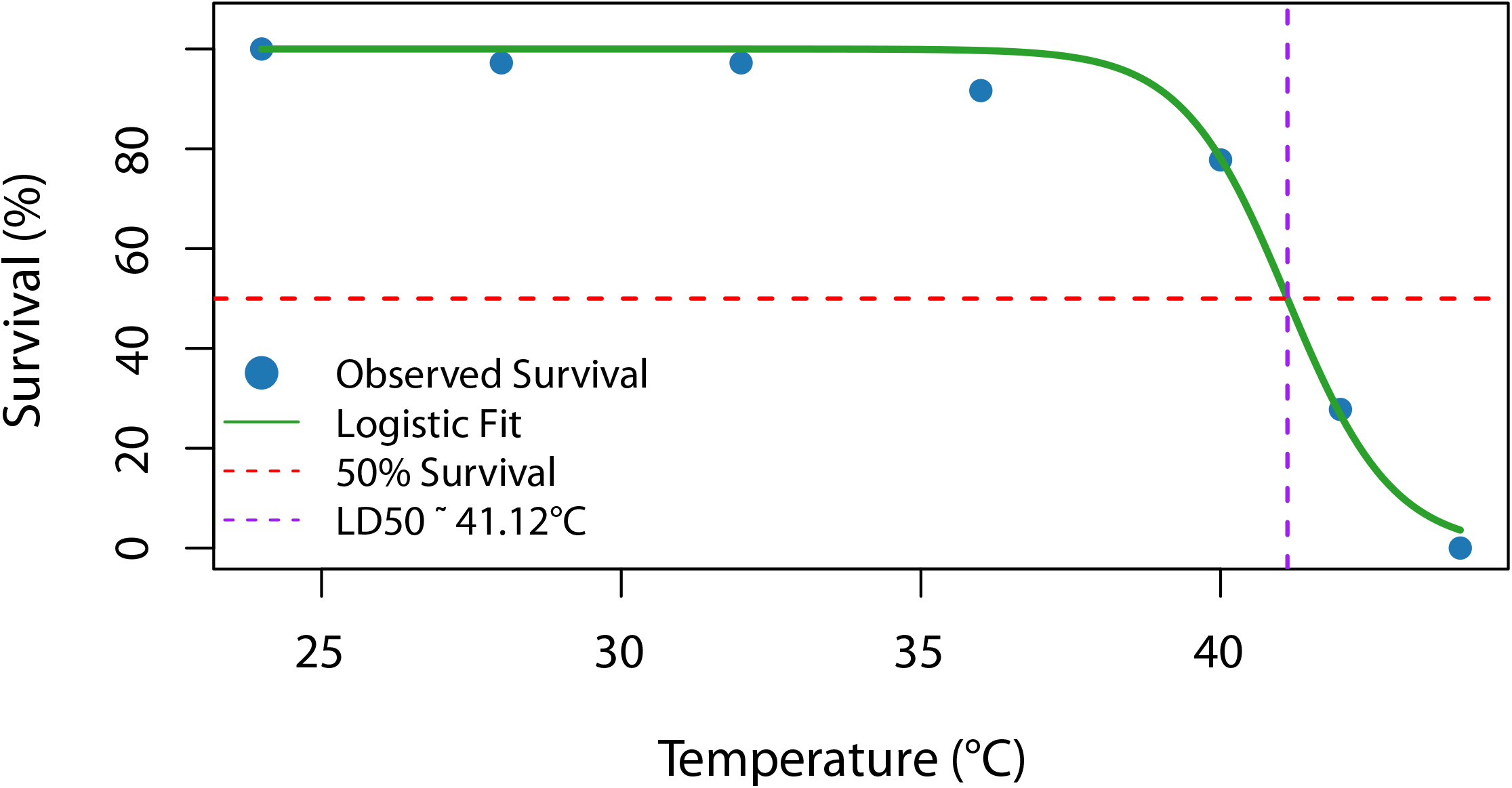
Survival of Rhodnius prolixus following acute heat stress. Fourth-instar nymphs (n=36 bugs per temperature point) were exposed to acute heat stress for 8 H across a range (24-44 °C). The blue circles represent observed mean survival (%), and the green line shows the logistic regression fit. The dashed red line indicates 50% survival, and the dashed purple line marks the estimated lethal dose for 50% mortality (LD_50_ = 41.1 °C). Error bars represent the standard error of the mean (SEM).

### Heat stress reduced blood feeding

Heat stress significantly affected the amount of blood ingested and the proportion of kissing bugs that ingested blood (Figures 3A and 3B). At one day post-exposure, bugs exposed to 24 °C consumed the largest blood meals (mean = 0.19 g, n = 36), while those exposed to 32 °C and 40 °C consumed progressively smaller meals (mean = 0.17 g and 0.11 g, respectively; n = 36). The difference in mean meal size across the three temperature groups was statistically significant (F(2, 15) = 23.90, p = 2.2 × 10^−5^). Pairwise comparison of bugs exposed to 40 °C fed significantly less blood than those at 24 °C (*p* = 8.5 × 10^−5^) and 32 °C (*p* = 0.00064) and feeding at 32 °C was reduced compared to 24 °C (*p* = 0.012). Proportion followed a similar trend, with all insects fed at 24 °C (0.97 ± 0.02) and 32 °C (0.94 ± 0.03), but only ∼75% fed after exposure to 40 °C. After 7 days of recovery, kissing bug feeding was restored across the groups. Meal size was comparable among treatments (24 °C: 0.20 g; 32 °C: 0.19 g; 40 °C: 0.17 g; all *p* > 0.25), and feeding proportion (Figure 3B) recovered to similar levels to those of controls (24 °C: 0.97; 32 °C: 0.94; 40 °C: 0.72). In summary, acute, sublethal heat stress reduced feeding after one day of recovery, but effects were diminished in bugs recovered for one week before allowing access to a bloodmeal.

**Figure 3.**
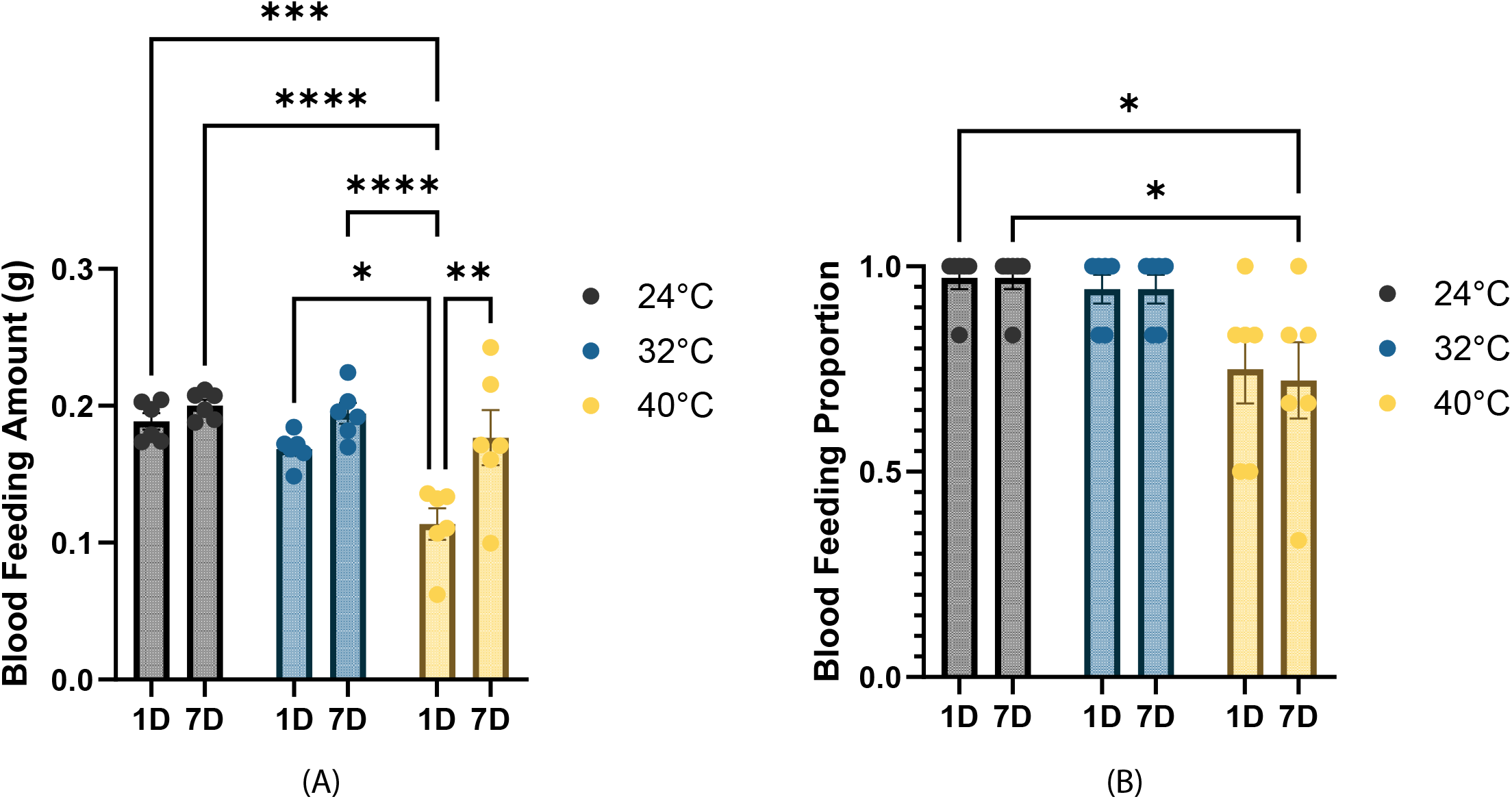
Effect of heat exposure on blood feeding amount and proportion after 1 day and 7 days of recovery for Rhodnius prolixus. **(A)** Blood feeding amount (g) of *R. prolixus* nymphs following an 8-h heat exposure at 24 °C, 32 °C, or 40 °C (n = 36 per temperature), measured after 1 day (1D) or 7 days (7D) of recovery. Data are shown as mean ± SEM. Feeding amount was analyzed using two-way ANOVA, revealing significant main effects of recovery time (F(1, 30) = 15.21, P = 0.0005) and temperature (F(2, 30) = 11.71, P = 0.0002), with no significant interaction between factors (F(2, 30) = 3.18, P = 0.056). Post hoc comparisons were performed using Tukey’s multiple comparisons test, with significant differences observed between 40 °C and both 24 °C and 32 °C at each recovery time, as well as between 1D and 7D within selected temperature groups (P < 0.05). **(B)** Blood feeding proportion of *Rhodnius prolixus* nymphs following an 8-h heat exposure at 24 °C, 32 °C, or 40 °C (n = 36 per temperature), measured after 1 day (1D) or 7 days (7D) of recovery. Bars represent mean ± SEM. Feeding success was analyzed using two-way ANOVA, which revealed a significant main effect of temperature (F(2, 30) = 10.20, P = 0.0004), but no significant effect of recovery time (F(1, 30) = 0.039, P = 0.844) and no interaction between temperature and recovery (F(2, 30) = 0.039, P = 0.961). Overall, feeding success remained high at 24 °C and 32 °C, whereas exposure to 40 °C significantly reduced the proportion of insects that successfully fed.

### Heat stress does not impact molting

Molting rates were comparable across all treatment groups, and neither temperature nor recovery duration produced statistically significant differences (two-way ANOVA; p > 0.05 for all factors) (Figure 4). Nymphs that did not experience heat exposure exhibited relatively high and consistent molting rates for both recovery periods, indicating normal post feeding development under control temperature. Molting success following exposure to 32 °C was slightly lower than that of those that did not experience thermal stress. Following exposure to 40 °C, nymphs displayed slightly lower molting than in the other temperature groups. However, these reductions were modest and not statistically significant. These findings indicate that short-term heat exposure has subtle, non-significant effects on molting success under the conditions tested.

**Figure 4.**
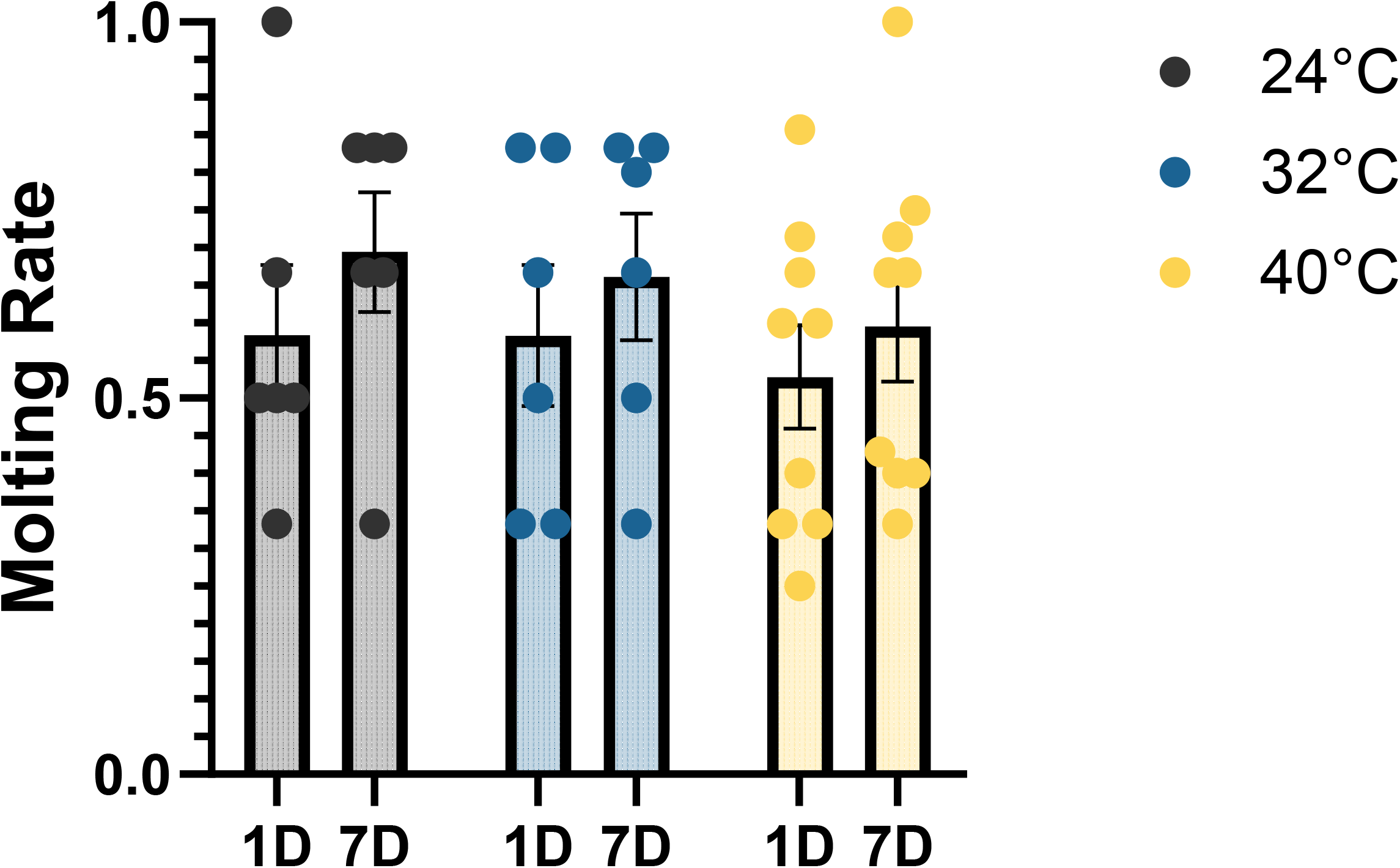
Molting rate of Rhodnius prolixus following heat exposure, recovery, and blood feeding. Molting rate of *R. prolixus* nymphs following an 8-h heat exposure at 24 °C, 32 °C, or 40 °C (n = 36 per temperature), measured after either 1 day (1D) or 7 days (7D) of recovery and subsequent blood feeding. Bars represent mean molting rate ± SEM. Molting success was analyzed using two-way ANOVA, which revealed no significant effects of temperature (F(2, 36) = 0.545, P = 0.585), recovery time (F(1, 36) = 1.603, P = 0.214), or their interaction (F(2, 36) = 0.0389, P = 0.962).

### Trypanosome infection is reduced following heat exposure

Heat stress significantly affects infection rates and blood feeding behavior in kissing bugs (Figures 5A and 5B). One day after exposure to 40 °C, infection levels decreased by nearly 60%. A less pronounced reduction was observed 7 days post-exposure, where bugs exposed to 40 °C exhibited lower infection rates than those at 24 °C, but this was no longer significant compared to the control. This suggests that recovery time partially restored feeding capacity and parasite acquisition. A reduction in bloodmeal size was previously observed and was examined in relation to infection status. To examine the observed reduction in bloodmeal size in relation to infection status, we found a significant interaction across all treatments (F(1, 104) = 21.52, p < 0.001). This interaction indicates that a reduction in bloodmeal size is associated with lower infection rates. This difference could explain the reduced infection rates observed when bugs were offered a bloodmeal one day after exposure to 40 °C, as over 60% of the bugs ingested a suboptimal bloodmeal (Figures 5A and 5B).

**Figure 5.**
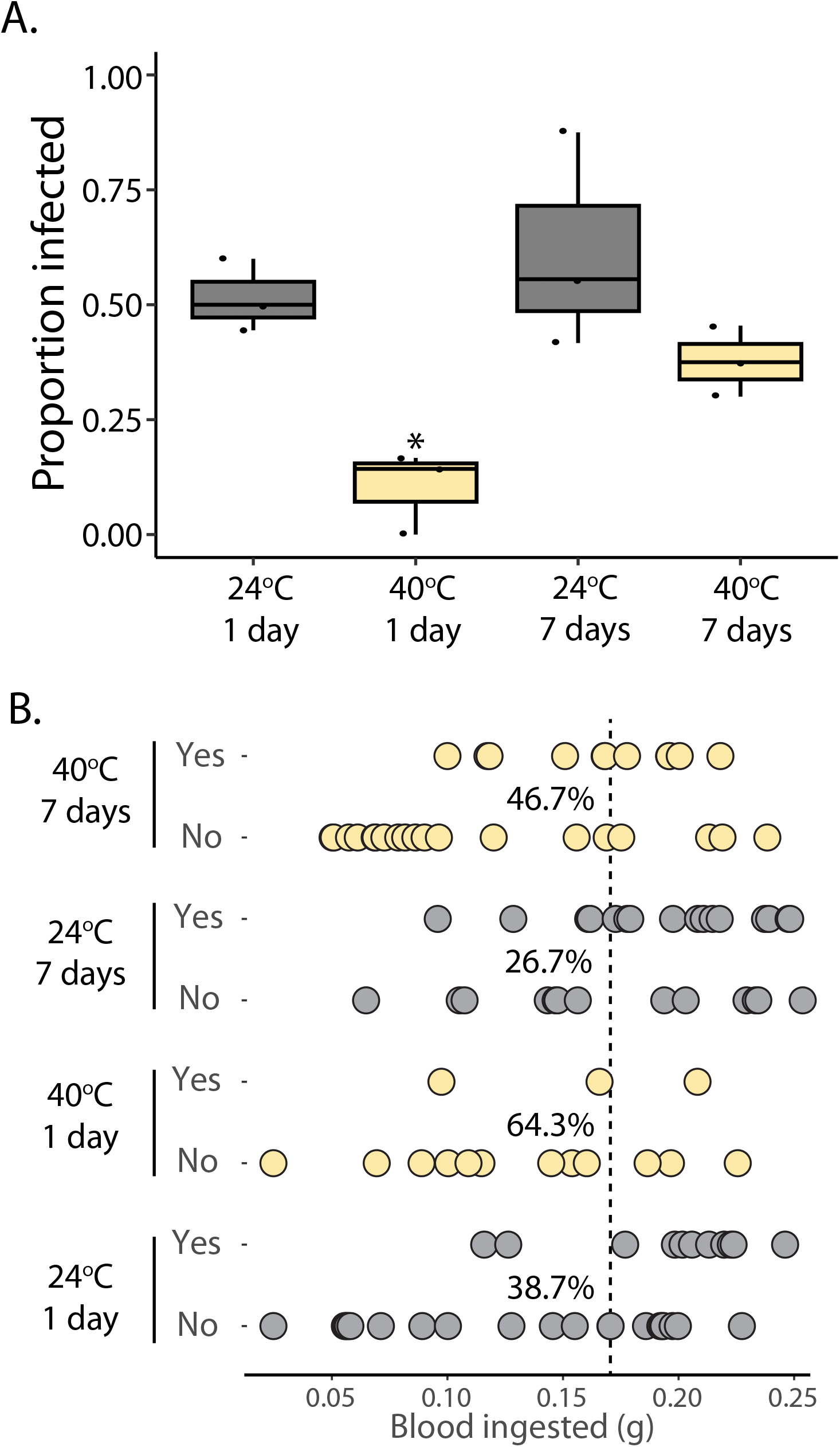
Infection rate of Rhodnius prolixus following heat exposure and recovery. **(A)** The infection rate of *R. prolixus* nymphs following an 8-hour heat exposure at 24 °C, 32 °C, or 40 °C (n = 36 per temperature), measured in the 1-day (1D) or 7-day (7D) recovery treatment group. The infection rate was significantly lower at 40 °C than at 24 °C after 1 day of recovery, with additional differences observed between 1D and 7D within some temperature groups. **(B)** The distribution shows that 40°C groups (both 1 day and 7 days) generally cluster lower along the x-axis compared to the 24°C controls, suggesting smaller bloodmeal sizes and potentially altered infection outcomes. Exposure to 40 °C reduced bloodmeal size relative to the 24 °C control. This reduction in feeding volume appeared to influence infection dynamics, as shown by altered distributions in infection status (Yes/No) across treatments.

## Discussion

Short-term heat stress significantly impacted the survival and blood feeding behavior of *R. prolixus*. Heat-exposed insects exhibited reduced survival and blood feeding rates compared with controls. A recovery period (one week) partially mitigated these effects, indicating a degree of physiological resilience following the high temperature exposure. Heat stress adversely affected parasite-associated outcomes, as infection rates declined following heat exposure, most likely due to reduced bloodmeal size. Short-term heat stress did not significantly alter molting success.

Previous studies have found that temperature affects the development of kissing bugs (da Silva and da Silva, 1989; Cabello, 1999; Luz et al., 1999; Mesquita et al., 2015). In this study, we found that acute heat stress significantly impacts the survival of kissing bugs. All individuals survived exposure at 24 °C, and only minimal mortality was observed at 32 °C, consistent with prior reports indicating that triatomines develop optimally within the mid 20 °C to low 30 °C range (Guarneri et al., 2003). In contrast, survival declined sharply following exposure to 40 °C, with complete mortality observed at 44 °C and an estimated LT_50_ of 41.1 °C. These results closely align with previous studies demonstrating a narrow upper thermal tolerance in triatomines, in which small increases above optimal temperatures lead to disproportionate mortality (Catalá et al., 2015; Belliard et al., 2019). The critical thermal temperatures of most insects range from 30 °C to 50 °C (Kellermann et al., 2012; Araújo et al., 2013). The reported critical thermal limits for bed bug survival (43.5–48 °C) (Benoit et al., 2009; Pereira et al., 2009; Kells and Goblirsch, 2011) are notably higher than those observed for *R. prolixus*, indicating that kissing bugs may be more vulnerable to acute heat exposure than other blood feeding hemipterans. Upper thermal tolerance parameters, including critical thermal maxima (CTmax), lethal temperature thresholds (LT_50_), and acute survival limits, are essential for predicting species survival, distribution, and responses to climate change (Terblanche et al., 2011; García-Robledo et al., 2016).

The amount of blood fed and the proportion of insects were lower following exposure to elevated temperatures, with the largest reductions observed after treatment at 40 °C compared with 24 °C when blood was offered one day after thermal exposure. Similar temperature-dependent declines in feeding rate have been reported in other triatomine species, including *Triatoma infestans* (Álvarez-Duhart et al., 2024) and *Triatoma brasiliensis* (Guarneri et al., 2003), suggesting a conserved response to thermal stress that reduces subsequent blood feeding. Comparable effects have also documented in mosquitoes and other vector-borne insects, in which high temperatures reduce the likelihood of blood feeding, including in *Aedes albopictus* (Muturi et al., 2012; Alto and Bettinardi, 2013). Notably, significantly fewer female mosquitoes fed on blood at 32 °C compared with 26 °C or 29 °C, indicating that even moderate thermal elevation can suppress feeding behavior. Importantly, blood feeding itself represents a significant thermal challenge: ingestion of a warm blood meal induces a protective heat-shock response in mosquitoes, including rapid upregulation of heat-shock proteins (Benoit et al., 2011; Lahondère et al., 2017). Together, these findings provide evidence that reduced feeding at elevated temperatures reflects the combined costs of ambient heat stress followed by thermal load imposed by the blood meal, which may constrain feeding success across diverse blood feeding vectors in relation to thermal stress. Molting in triatomines is constrained by a critical period of approximately 7 days following blood feeding, and chronic exposure to elevated temperatures can delay or suppress molting by sustaining metabolic disruption (Wigglesworth, 1952). In contrast, the present study focuses on short-term, acute heat stress, and we observed no significant differences in molting success under any recovery condition. This indicates that brief thermal exposure, even when it alters post-stress physiology, is insufficient to disrupt the molting program, as long as the kissing bug ingests a bloodmeal. Instead, our results demonstrate that acute thermal stress primarily affects short-term post-feeding performance, and that the timing of recovery modulates these effects without crossing the threshold required to impair molting.

Previous studies have shown that temperature strongly influences *T. cruzi* development in triatomine vectors, particularly under prolonged thermal exposure. In *R. prolixus*, infection rates increase with temperature within a sub-extreme range, with temperatures between 20 °C and 34 °C promoting parasite establishment and replication, and parasite loads peaking near 28 °C (Elliot et al., 2015; Loshouarn and Guarneri, 2024). Similarly, González-Rete et al. (González-Rete et al., 2021) reported that chronic exposure of *Triatoma pallidipennis* to 34 °C resulted in lower parasite loads than at 20 °C or 30 °C, indicating that sustained high temperatures can reduce parasite establishment in kissing bugs. Importantly, these studies involved continuous exposure for days to weeks, either before or throughout parasite development. Our study tested the effects of a short, sub-lethal heat exposure applied immediately prior to blood feeding, followed by recovery under standard conditions. Using two discrete temperature treatments (24 °C and 40 °C), we found that acute heat stress at 40 °C significantly reduced parasite burden in *R. prolixus*. This reduction was accompanied by a marked decline in blood feeding rate at 40 °C, which was associated with parasite presence, suggesting that acute thermal stress prior to feeding indirectly limits parasite establishment by impairing feeding. Limited blood intake, which reduces trypanosome infection, is consistent with previous studies demonstrating that blood meal size is a key determinant of parasite establishment in triatomines (Asin and Catalá, 1995). Together, these findings indicate that the timing and duration of thermal exposure are critical determinants of infection outcomes, with acute pre-feeding heat stress effects that are distinct from those observed under prolonged thermal conditions.

Although parasites were not directly exposed to elevated temperatures, acute heat stress prior to feeding may have indirectly influenced parasite establishment by altering the host midgut environment during the early post-feeding period, a critical period for *T. cruzi* survival and differentiation within triatomines (Kollien and Schaub, 2000). More broadly, vector competence is highly sensitive to host physiological state and environmental conditions, particularly under sustained exposure during parasite acquisition or development (Kollien and Schaub, 2000; Thomas and Blanford, 2003). Together, acute thermal stress experienced prior to feeding could indirectly influence infection outcomes by affecting host feeding performance and nutrient availability, rather than through direct thermal effects on the parasite.

Our study highlights the importance of pre-feeding host condition following acute thermal stress as a previously underappreciated determinant of vector competence (Kobayashi et al., 1981; Arthurs and Thomas, 2000; Thomas and Blanford, 2003). These results align with broader insect physiology studies showing that recovery from sublethal stress occurs over days and can restore feeding and developmental capacity; here, we extend that concept to parasite acquisition and vector competence in kissing bugs in relation to thermal exposure. Specifically, this study defines the lethal temperature threshold for *R. prolixus* under acute thermal stress and demonstrates that short-term heat exposure produces a suite of sublethal effects on key traits relevant to vector performance. Importantly, we show that these sublethal effects are strongly modulated by recovery time prior to feeding, with blood meal size, feeding success, and parasite establishment largely recovering when sufficient time is allowed between heat exposure and blood ingestion. As climate change is expected to increase the frequency and intensity of heat waves (Perkins-Kirkpatrick and Lewis, 2020), short-term exposure to extreme temperatures may pose a significant challenge to vector persistence and influence disease transmission dynamics. Overall, these findings provide critical insights into how extreme thermal events could impact kissing bug biology and reduce vector competence for *T. cruzi*.

## ACKNOWLEDGMENTS

Partial funding for reusable equipment was provided indirectly by the National Institute of Allergy and Infectious Diseases, R01AI148551 and R21AI166633 to J.B.B. Partial funding for parasite studies was provided by the National Institute of Allergy and Infectious Diseases of the National Institutes of Health (R00AI137322 and R21AI182544 to N. Lander). The funding agencies had no role in the study design, data collection, interpretation, or the decision to submit the work for publication. Opinions contained in this publication do not reflect the views of the funding agencies. We declare that we have no competing financial interests.

